# The cancer-mutation network and the number and specificity of driver mutations

**DOI:** 10.1101/237016

**Authors:** Jaime Iranzo, Iñigo Martincorena, Eugene V. Koonin

## Abstract

Cancer genomics has produced extensive information on cancer-associated genes but the number and specificity of cancer driver mutations remains a matter of debate. We constructed a bipartite network in which 7665 tumors from 30 cancer types are connected via shared mutations in 198 previously identified cancer-associated genes. We show that 27% of the tumors can be assigned to statistically supported modules, most of which encompass 1-2 cancer types. The rest of the tumors belong to a diffuse network component suggesting lower gene-specificity of driver mutations. Linear regression of the mutational loads in cancer-associated genes was used to estimate the number of drivers required for the onset of different cancers. The mean number of drivers is ~2, with a range of 1 to 5. Cancers that are associated to modules had more drivers than those from the diffuse network component, suggesting that unidentified and/or interchangeable drivers exist in the latter.

## Introduction

Cancer develops as a result of accumulation of somatic mutations that impair cell division checkpoints, resulting in abnormal cell proliferation and eventually tumorigenesis^1, 2^. Such mutations are called “drivers” because they are thought to drive their carrier towards a cancerous state. Characterization of driver mutations is central to understanding the early steps of tumor progression^3, 4^. During the last decade, comparative analyses of large collections of cancer genomes have led to the identification of overlapping sets of genes that are typically associated to cancer, i.e. harbor a significant excess of mutations in tumors and show signatures of positive selection^5–10^. The continued identification of cancer-associated genes provides insights into the processes and pathways involved in tumorigenesis, as well as possible therapy targets^11–16^. Like any other gene in the genome, cancer-associated genes are expected to accumulate passenger mutations that do not contribute to or even hinder cancer progression^17^. Therefore, although cancer-associated genes harbor numerous driver mutations, only a fraction of the mutations found in these genes are actual drivers^10^.

Distinguishing driver mutations from passenger ones poses a formidable challenge for cancer genomics. The number of driver mutations required for the onset of cancer is a fundamental question that remains a matter of debate^3, 9, 18–20^. Classical approaches to this problem use age incidence curves to infer the number of rate-limiting steps in tumorigenesis, each of which is assumed to be associated with a unique driver mutation; these estimates, however, are sensitive to changes in mutation and replication rates during tumor progression^19–22^. A modified method has been proposed that compares the incidence of cancer across risk groups with different mutation rates, but this approach applies only to cancers with relatively well-defined risk groups, such as lung and colorectal cancer^18^. Recent measurements of selection in cancer genomes have provided quantitative estimates of the number of positively selected mutations, i.e. drivers, per tumor ranging from <1 in thyroid and testicular cancers to >10 in endometrial and colorectal cancers^9^. Given the novelty of these findings, a comparison with independent inference approaches appears highly desirable.

A second major question about cancer driver mutations refers to their specificity in different cancer types. Some tumors show recurrent mutation patterns, such as the oncogenic fusion BCR-ABL in chronic myeloid leukemia23 or the inactivation of specific tumor suppressors such as, for example, RB1 in retinoblastoma^24^. Other tumors appear to result from interchangeable mutations in a pool of genes involved in key signaling pathways, such as the receptor tyrosine kinase/RAS/RAF pathway in lung adenocarcinoma^25^. Between these two extremes, intermediate degrees of specificity are observed in many cancer types^8, 26, 27^. Furthermore, although numerous recent studies on cancer mutational landscapes have yielded extensive lists of genes that are mutated in various cancers^28–30^, a quantitative understanding of the extent to which the current tumor classification captures the existence of specific sets of driver mutations is lacking.

Here, we combine tools for network analysis and multivariate statistics to assess the number and specificity of cancer driver mutations in 30 cancer types. We show that an unsupervised community detection approach applied to the bipartite network of somatic mutations in cancer recovers modules consisting of mutually specific tumors and genes that (i) are consistent with the tumor histology and (ii) are enriched in putative driver mutations. We used multivariate statistical analysis to estimate the characteristic number of driver mutations in cancer-associated genes required for the onset of each cancer type. Notably, the average age of onset for different cancer types correlates with the predicted number of drivers. Furthermore, cancers that are not associated with the specific modules in the gene-tumor network appear later in life than expected based on the general trend.

## Results

### Cancer mutational landscape as a partially modular network

Somatic mutations in a set of tumors can be collectively represented as a bipartite network, that is, a network with two classes of nodes. In such a network, nodes of one class correspond to tumor samples and nodes of the other class correspond to cancer-associated genes. Mutations are represented as edges that connect each tumor sample with the genes mutated in it; conversely, each gene is linked to the tumors in which it carries a mutation(s). Using this approach, we built the network of somatic mutations from The Cancer Genome Atlas (TCGA), a collection that consists of 7665 tumor samples from 30 cancer types (Fig. 1A), focusing on coding mutations in 198 recurrently mutated cancer genes (see Methods).

**Figure 1:**
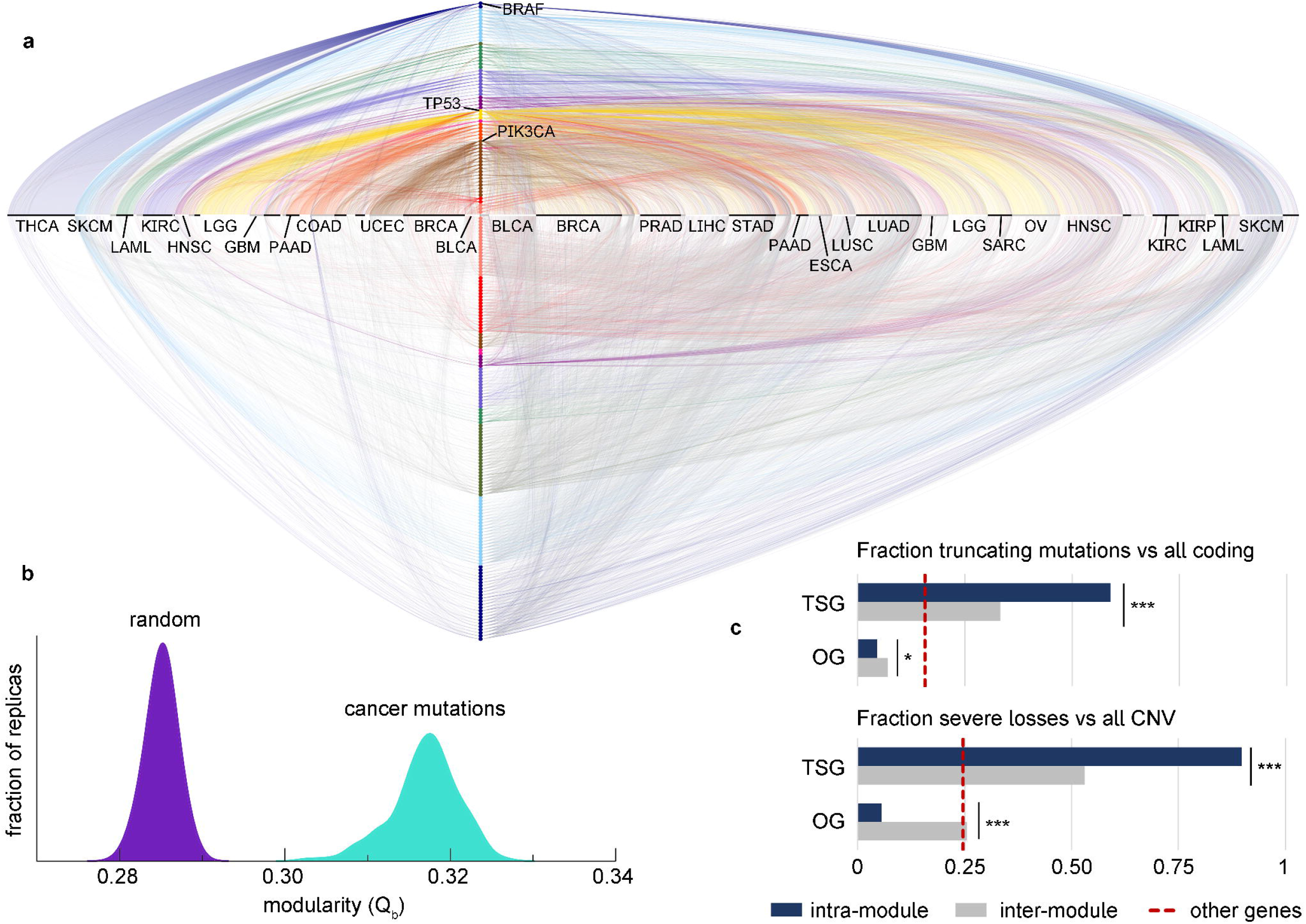
Modular structure of the cancer mutation network. **(a)** Bipartite network of somatic mutations in tumors from the TGCA. Samples are arranged by cancer type along the x axis; cancer-associated genes are sorted by module along the y axis. Samples from the same cancer type and genes from the same module are sorted by degree. The upper and left semi-axes contain genes and samples that belong to statistically significant modules. The rest of nodes (lower and right semi-axes) were assigned to the “best-match” extended module with which they share the highest similarity (see text). Links connect samples and genes affected by at least one nonsynonymous somatic mutation. Links between two nodes from the same module (intra-module links) are drawn in distinctive colors; inter-module links appear in gray. Cancer type abbreviations are given in Supplementary Table S1. **(b)** The modularity of the whole cancer mutation network was quantified by its Barber’s modularity index (Qb) and compared to 200 random networks with the same degree distribution. The modularity distribution for the cancer mutation network results from 200 realizations of the community detection algorithm, each yielding slightly different sets of modules. The lack of overlap reveals a highly significant (p<10^−20^, Welch’s T-test) modular structure for the cancer mutation network. **(c)** Differences in the functional spectrum of mutations between intra-module and inter-module links (significant modules only). Among small, coding mutations, truncating mutations typically constitute intra-module links in tumor suppressor genes (TSG) and inter-module links in oncogenes (OG). Among copy number variants, severe losses are typically observed among TSG and samples from the same module, whereas OG losses are typically observed in samples that belong to a different module. Asterisk indicate the level of statistical significance (* p<0.05, *** p<10^−5^).

Within the bipartite network framework, the association between mutually specific sets of genes and cancer types becomes manifest by the existence of groups of nodes (tumors and genes) that are densely connected with members of the same group but poorly connected with the rest of the network. Such groups are called “modules”, and a network with such structure is said to be modular^31^. We tested the modular nature of the cancer mutation network by computing its Barber’s modularity index32 and comparing it with 200 random networks with the same degree distribution (Fig. 1B). The result of this comparison supported the existence of a significant degree of mutual specificity between cancer types and cancer-associated genes (p<10^−20^, Welch’s T-test), which demonstrates the ability of the network approach to detect a well known feature of cancer mutational landscapes^3, 8, 26^, To further investigate the specificity of the mutation landscapes, we identified the modules of the network and assessed their statistical significance. To that end, we first applied a battery of module detection algorithms and then pruned all genes and tumors for which the specificity patterns were compatible with a random null model (see Methods). The analysis revealed the existence of 12 modules with a significance threshold p < 0.05. Each of these modules contains tumors and genes that are mutually specific, i.e. tumors in a module typically harbor mutations in genes from the same module, whereas the constituent genes are more frequently mutated in tumors that belong to the same module. Overall, the statistically significant modules comprise 27% of the samples and 66 (33%) cancer associated genes (Table 1).

**Table 1:**
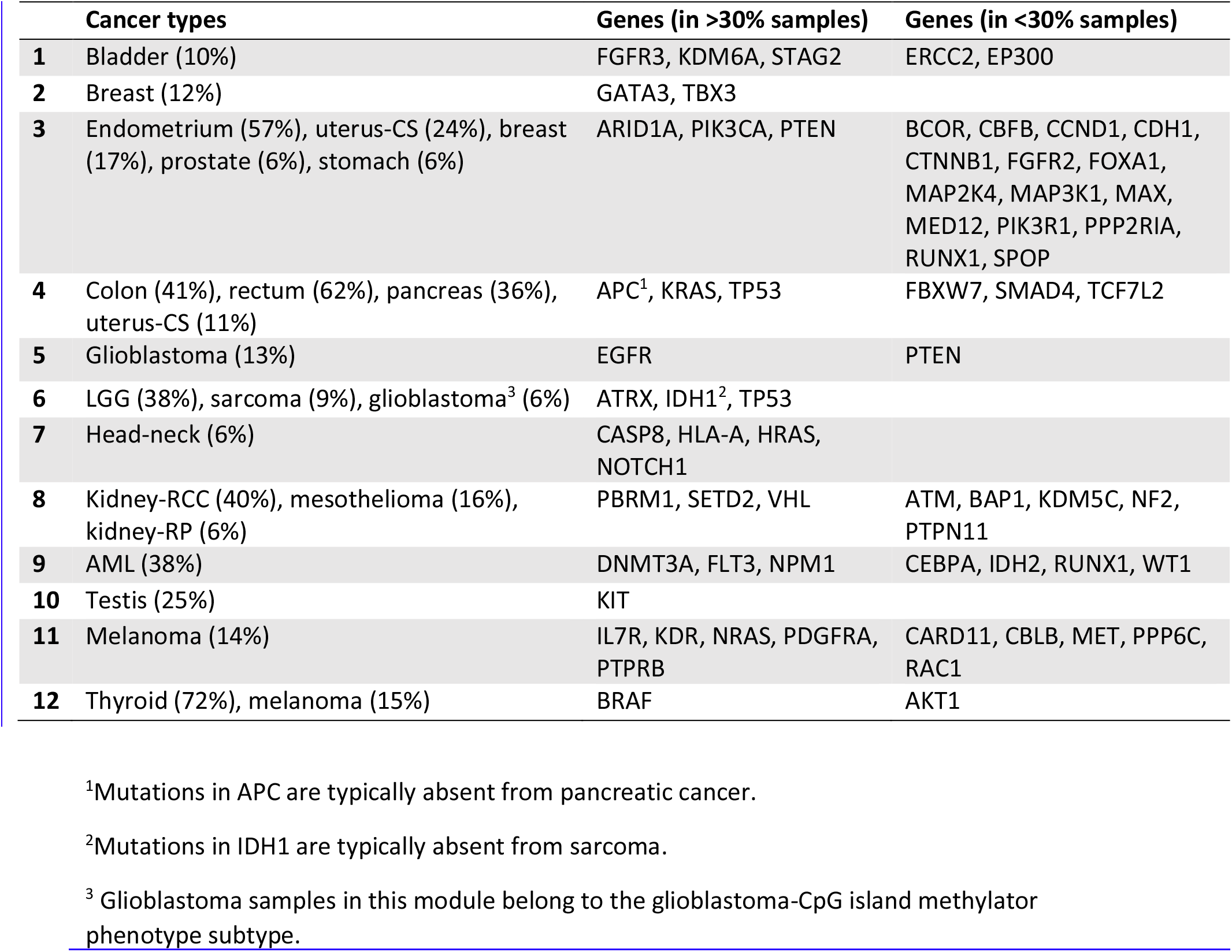
Composition of the 12 statistically significant modules in the cancer mutation network. Samples are grouped by cancer type; the number in parentheses indicates the fraction of samples from that class that are present in the module (only classes represented by >5% of their samples are shown). Kidney-RCC: kidney renal clear cell carcinoma, kidney-RP: Kidney renal papillary cell carcinoma, AML: acute myeloid leukemia, LGG: brain lower grade glioma, uterus-CS: Uterine Carcinosarcoma.

**Table 2:** Modules in the subnetwork of breast, prostate, endometrial and uterine cancers. Numbers indicate the percentage of samples from a given cancer type associated to the module.

| **Genes** | **Breast**<br>**(%)** | **Prostate**<br>**(%)** | **Endometrium**<br>**(%)** | **Uterus-CS**<br>**(%)** |
| --- | --- | --- | --- | --- |
| CBFB, CDH1, GATA3, MAP2K4,<br>MAP3K1, PIK3CA, RUNX1,<br>TBL1XR1, TBX3 | 30 | 2 | 2 | 0 |
| ARID1A, BCOR, CCND1, CIC,<br>CTNNB1, CUX1, ESR1, FGFR2,<br>KRAS, MAX, PIK3CA, PIK3R1, PTEN | 1 | 1 | 43 | 7 |
| FOXA1, SPOP | 3 | 26 | 2 | 5 |
| FBXW7, PPP2RIA, TP53 | 11 | 6 | 15 | 55 |

Before proceeding with a more detailed dissection of the genes and cancer types represented in each module, we evaluated their biological relevance by characterizing the mutations that correspond to intra- and inter-module connections. To that end, we split the cancer associated genes into oncogenes and tumor suppressor genes (TSG), and calculated the fraction of truncating mutations (with respect to all small nonsynonymous mutations) and deletions (with respect to all copy number variants) in each of these genes, in samples assigned to the same module as the gene (intra-module links), and in samples assigned to a different module (intermodule links). We found that intra-module connections include a significantly greater fraction of truncating mutations in TSG than inter-module connections, whereas the opposite holds for oncogenes (Fig. 1C). A similar trend is observed in copy number variation data: intra-module connections encompass a significantly higher fraction of TSG loss and oncogene amplification compared to inter-module connections. Overall, intra-module connections are significantly enriched in putative driver mutations including truncating mutations and gene loss in TSG, and missense mutations and amplification in oncogenes. Thus, mutations that affect mutually specific genes and tumors (i.e. genes and samples from the same module) are more likely to be cancer drivers than those affecting genes and samples that belong to different modules. Nevertheless, deviations with respect to the baseline fraction of truncating mutations in noncancer-associated genes indicate that some mutations involving genes and tumors from different modules are also relevant for tumor progression. Such deviations remain after removing the most widespread cancer genes across tissues (TP53, PIK3CA, and ARID1A), indicating that potential inter-module drivers are not limited to such genes (Supplementary Fig. S1). A closer inspection of inter-module mutations highlights oncogenes BRAF and IDH1 as major sources of inter-module driver mutations in melanomas and acute myeloid leukemia, respectively, with missense mutations representing 96% of the coding mutations in both genes compared to the 84% baseline. Similarly, tumor suppressor genes STAG2, KDM6A, PIK3R1, MAP3K1, and CDH1, with 50-60% of truncating mutations (16% baseline), constitute probable inter-module drivers, with MAP3K1 and CDH1 being more relevant in breast cancer.

### Specificity modules for cancer types and cancer-associated genes

As shown in Table 1, the composition of the gene-sample specificity modules strongly correlates with the histological classification of tumors. Most modules include tumors from one or two cancer types, together with genes for which mutation frequencies are significantly higher in cancers of those types (Fig. 2). The mutual specificity analysis recovers some well-established features of cancer mutational landscapes, such as the association between thyroid cancer and BRAF, or between colorectal cancer and APC, KRAS, TP53, and SMAD4. The colorectal cancer module includes two additional genes that are mutated in a smaller fraction of samples, namely, ubiquitin ligase FBXW7 and transcription factor TCF7L2. Most samples from pancreatic adenocarcinoma cluster in the same module as colorectal cancer, in agreement with a significant excess of mutations in KRAS, TP53 and SMAD4 in both tumor types. Acute myeloid leukemia constitutes a single module, with the genes DNMT3A, FLT3 and NPM1 mutated in >30% samples, and CEBPA, IDH2, RUNX1 and WT1 mutated at lower frequencies. Testicular germ cell cancer, head and neck squamous cell carcinoma, and urothelial bladder carcinoma also form separate modules which, however, comprise a smaller fraction of the samples from each cancer type. The sets of associated genes include KIT in testicular cancer; NOTCH1, CASP8, HLA-A and HRAS in head and neck cancer; and FGFR3, KDM6A and STAG2 in bladder cancer. Clear cell kidney carcinoma clusters with genes VHL, PBRM1, SETD2 and BAP1, among others. Some samples from mesothelioma and papillary kidney carcinoma are also assigned to that module, mostly because of mutations in SETD2 and BAP1.

**Figure 2:**
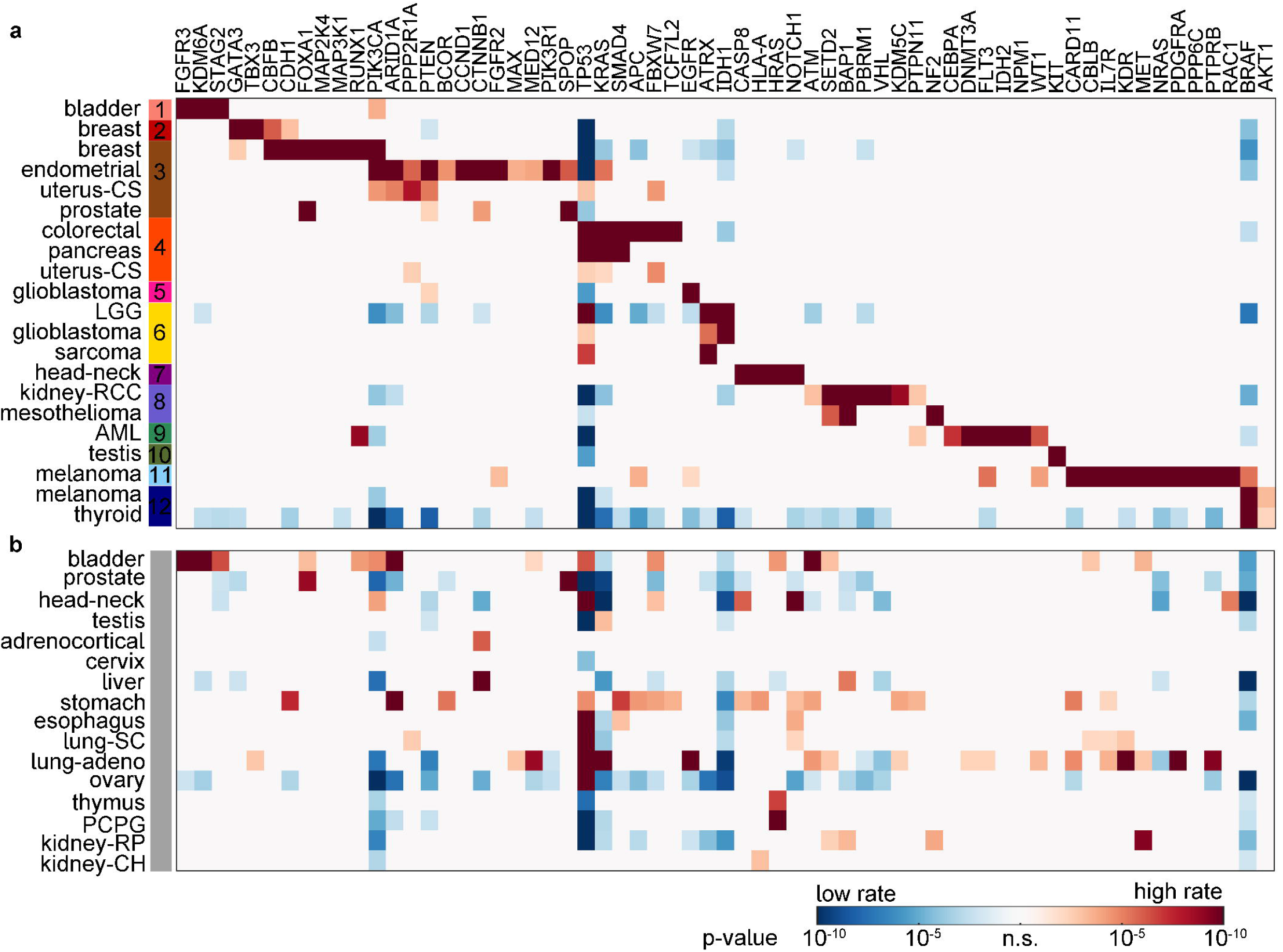
Cancer genes mutated at significantly distinct rates in different modules and cancer types. Tumors that do and do not belong to specificity modules are shown in **(a)** and **(b)**, respectively. Only genes that belong to specificity modules are shown. Significance was evaluated with a two-tailed Fisher’s exact test; red (blue) indicates a higher (lower) than average prevalence of mutations.

Notably, some cancer types are distributed among more than one module. Thus, glioblastoma is represented in two modules, one characterized by mutations in EGFR and PTEN, and the other by mutations in IDH1, ATRX and TP53. Such subdivision is consistent with previous reports, which relate the second group with the glioblastoma-CpG island methylator phenotype^33–35^. The same module also includes most samples from lower grade glioma and some representatives from sarcoma (though the latter typically lack mutations in IDH1). Similarly, melanoma is divided into tumors with mutated BRAF on a low mutational background, which cluster with thyroid cancer, and samples with mutations in a larger set of genes (including NRAS, KDR, ILR7, PTPRB), which constitute a separate module. Finally, breast cancer splits between a breast cancer-only module characterized by mutations in GATA3 and TBX3, and a larger module that includes uterine (endometrial and carcinosarcoma) and prostate cancers, with PIK3CA as the signature gene. In terms of histological types, the PIK3CA module is significantly enriched in lobular breast tumors (46% compared to 13% in the GATA3/TBX3 module and 17% in the entire dataset, Chi-squared test *p* < 10^−6^). In terms of the molecular subtypes, both modules include breast tumors that mostly belong to the luminal subtype (estrogen receptor-positive), whereas most of the basal-like breast tumors are not assigned to any significant module (Chi-squared test *p* < 10^−10^).

Among all the modules, the one that combines breast, uterine and prostate cancers stands out for its size and diversity. This module contains the largest number of genes, with many of those mutated in less than 30% of the samples. Moreover, two of its constituent histologies, breast cancer and uterine carcinosarcoma, are split between this and other modules. To further dissect the specificity of the mutations affecting these cancer types, we reanalyzed the subnetwork composed by all cancer genes and samples from breast, prostate and uterine (endometrial and carcinosarcoma) cancers. This analysis yielded 4 significant modules that are dominated by each of the 4 cancer types. The list of module-specific genes is consistent with the findings of the global analysis. Notably, the re-analysis places most breast cancer samples in a single module with genes CDH1, PIK3CA, GATA3 and TBX3, whereas uterine carcinosarcoma clusters with genes FBXW7, PPP2RIA and TP53 (the samples without mutations in PPP2RIA were formerly assigned to the colorectal cancer module). Specific modules for prostate and endometrial cancer are also clearly delineated, the former with SPOP and FOXA1, the latter with ARID1A, CTNNB1, PI3KR1 and PTEN, among other genes.

### Two major modes of driver accumulation

The statistically significant modules of mutual tumor-gene specificity include 27% of the tumors in the TGCA. There are at least two alternative explanations for why 73% of samples remain unassigned. The first possibility is that, despite having mutational patterns compatible with one of the modules, the unassigned samples do not reach the required threshold of statistical significance. That would be the case if mutations affecting module-specific genes occurred in non-coding regions, involved copy number variants, or else, if mutations occurred in functionally equivalent genes not included in our list of cancer-associated genes. The second possibility is that unassigned samples account for exchangeability of cancer-associated genes. Under such a scenario, some cancer types might not be specifically associated to any set of genes.

To evaluate the first possibility, we built a set of “best-match extended” modules by attaching unassigned samples and genes to the module with which they shared most connections (Supplementary Table S2). We would expect that, if the specific association between tumors and genes held for most samples within a cancer type, the extended modules would recover unassigned samples from the same cancer types as those already assigned to the original modules. Indeed, the best-match extended modules cluster >75% of the samples from rectum, pancreas, kidney (clear cell), acute myeloid leukemia, thyroid and melanoma, and 50-75% of the samples from lower grade glioma, mesothelioma, colon and endometrial cancer in a tissue-specific way (Fig. 3A). In contrast, cancer types that were absent from the original modules appear distributed among multiple extended modules, with typically <25% of the samples assigned to the same module (Fig. 3B). The only exception is ovarian cancer, which does not appear in any of the significant modules although 70% of the samples are recovered as members of the same extended module as lower grade glioma. The apparent cause is the tight association between ovarian cancer and TP53, which is mutated in almost 90% of samples, against a low background of somatic mutations^36^.

**Figure 3:**
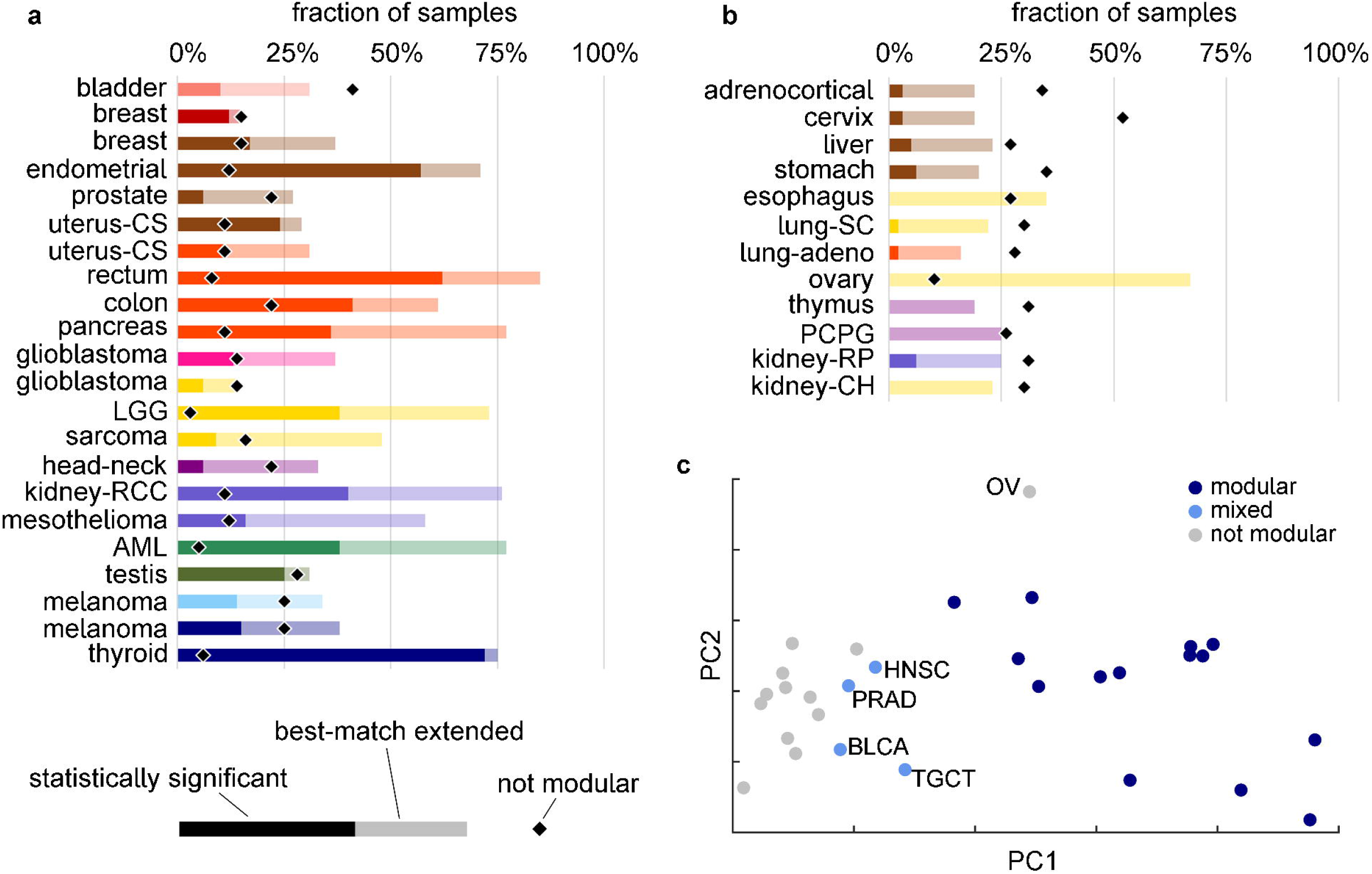
Classification of cancer types according to the gene-specificity of their driver mutations. **(a,b)** Fraction of samples assigned to statistically significant (solid bars) and best-match extended modules (semi-transparent bars), obtained by reassigning non-significant samples and genes to the significant modules with which they share the largest number of connections. The black diamond symbol indicates the fraction of samples assigned to the largest non-significant pseudo-module. Cancer types without major contributions to any significant module are shown in **(b)**; bar colors refer to the best-match extended module that contains most samples from each type. **(c)** Principal component analysis of cancer types based on the fraction of samples assigned to statistically significant modules, best-match extended modules, and the largest non-significant pseudo-module. The percentages of the total variance explained by the first and second components are 88.5% and 8.6%, respectively. Special cases discussed in the text are labeled: OV, ovarian; HNSC, head-neck; PRAD, prostate; BLCA, bladder; TGCT, testis.

We confirmed the existence of a diffuse (non-specific) mode of driver accumulation by running the module detection pipeline without removal of non-significant members. The resulting set of “statistically relaxed” modules consists of 12 modules with a counterpart among the original (statistically significant) modules, 16 minor modules with a single gene each and small sample sizes, and a giant, non-significant module that includes 20% of the samples and 90 (45%) genes (Supplementary Table S3). The appearance of non-significant (pseudo-)modules is a well-known artifact that results from applying module detection algorithms to large networks with partial or no modular structure^37, 38^. In the context of cancers and the associated genes, the giant pseudo-module accounts for the non-modular (diffuse) component of the cancer mutation network, which comprises exchangeable cancer genes that are not specifically associated to particular groups of tumors. Cancer types differ with respect to their contribution to this component. Thus, only 10-20% of samples from cancer types that are represented in the original modules are assigned to the diffuse component, whereas the fraction rises to more than 25% in other cancers (p < 0.01, Wilcoxon test; Fig. 3A and B).

Taken together, these findings indicate that cancer types can be conceptually split into two major groups: (i) those that accumulate driver mutations in specific sets of cancer genes and are accordingly clustered into distinct modules (Table 1), and (ii) those that accumulate exchangeable driver mutations in a non-tissue-specific manner, such as stomach and lung cancers. Although most cancer types can be clearly placed in one of those two extreme categories, there are some mixed cases (Fig. 3C). For example, bladder cancer generates its own module that includes genes KDM6A, FGFR3 and STAG2. However, only 10% of bladder tumors are assigned to that module (30% in the best-match extension), whereas about 40% belong to the diffuse component. Similar, albeit less extreme, trends are observed for head and neck and testicular cancers. Such a mixed mutational landscape, modular for some subsets of the samples, but non-specific for others, seems to mirror the heterogeneity of these cancer types.

### Copy number alterations

We further explored the specificity of driver events by jointly considering somatic mutations and copy number alterations that affect cancer-associated genes. To reduce the number of non-informative connections, only amplifications of oncogenes and losses of tumor suppressor genes were added to the network of somatic mutations. The addition of copy number alterations does not significantly change the results described so far (Supplementary Tables S4 and S5). The same significant modules are recovered, although their composition in terms of cancer types is slightly less clean. Besides the modules described above, the addition of the copy number alterations resulted in 5 new modules, none of which showed an obvious correspondence with a particular cancer type. Four of these modules are associated with arm-level alterations affecting chromosome arms 7q (genes BRAF, EZH2, MET, and SMO), 9q (genes ABL1, GNAQ, and KLF4), and 17p (genes MAP2K4 and NCOR1), and X chromosome region Xp11 (genes BCOR, GATA1, KDM5C, and KDM6A). The fifth of these new modules accounts for frequent loss of 56 cancer-associated genes (mostly tumor suppressors) distributed across the genome. Overall, the inclusion of oncogene amplifications and TSG losses does not reveal the existence of specific modules that were not already detected by the analysis of somatic mutations, although this outcome could be biased by the fact that most cancer-associated genes in our study were identified through mutations. The minor changes in the network structure caused by the inclusion of copy number alterations seem related to the chromosomal location of cancer genes and the opposite trends towards amplification and deletion in oncogenes and TSG, respectively.

### Average number of driver mutations

Identification of driver mutations is confounded by the numerous passenger mutations that are typically found in cancer genomes. Passenger mutations in the coding regions appear to dominate even in cancer-associated genes^9, 39^, resulting in a strong correlation between the number of mutations in cancer-associated and other genes (R = 0.87, p < 10^−20^, Supplementary Fig. S2). To obtain an estimate of the number of driver mutations in different tumors, we built a general linear model, with the number of coding mutations (substitutions and small indels) in cancer-associated genes as the dependent variable, the number of coding mutations in putative passenger genes as explanatory variable, and the cancer type as grouping factor. Due to the pervasive abundance of passenger mutations, a major feature of the model is the strong correlation between mutations in cancer-associated and non-cancer-associated genes. In this context, the intercepts (which depend on the cancer type) can be interpreted as the excess of mutations in cancer-associated genes that is not attributable to the same causes that lead to the accumulation of mutations in non-cancer-associated genes. Thus, these intercepts constitute a proxy for the number of driver mutations in cancer-associated genes.

We found that 75% of the variance in the number of mutations in cancer-associated genes is explained by the number of mutations in passenger genes (ANCOVA, p < 10^−20^), which is indicative of a common trend of mutation accumulation in both gene classes (Fig. 4A). Considering all tumors together and controlling for the non-uniform abundances of different cancer types, the mean number of driver mutations per tumor was estimated as 1.82 ± 0.07 (95% confidence interval). Differences in the intercepts across tumor classes are statistically significant but explain only 4% of the total variability in the data (Fig. 4A and B, Supplementary Table S6). Such low explanatory power could be due to the heterogeneity of the samples within the same cancer type and possible occurrence of driver mutations in non-coding regions or in genes that do not belong to our list of 198 cancer-associated genes; indeed, 12% of the samples lack coding mutations in these genes. Both limitations can be mitigated by analyzing only samples from significant modules, which represent more homogeneous subsets of tumors enriched in putative driver mutations. Thus, to assess how the number of driver mutations varies across tumors, we repeated the regression analysis with the samples that were assigned to significant modules, using both the assignment to the modules and the cancer type as the grouping variables (Fig. 4A, Supplementary Table S7). In samples from significant modules, differences in the number of drivers across cancer types explain 20% of the observed variance in the number of mutations in cancer-associated genes. The predicted number of driver mutations in these samples ranges from values near 1 in glioblastoma, thyroid carcinoma and the subset of melanoma with low mutational load, to values around 5 in bladder, endometrial and head and neck cancers (Fig. 4C). The average number of drivers per tumor in colorectal cancer is 3.66 ± 0.27, consistent with previous estimates based on epidemiological models^18, 40^ and measures of positive selection in cancer-associated genes^9^. Notably, the estimates of the average number of driver mutations in the two groups of melanomas differ by almost 2 (2.55±0.58 vs 0.68±0.39, respectively), which could be related to the higher mutation burden observed in the first group.

**Figure 4:**
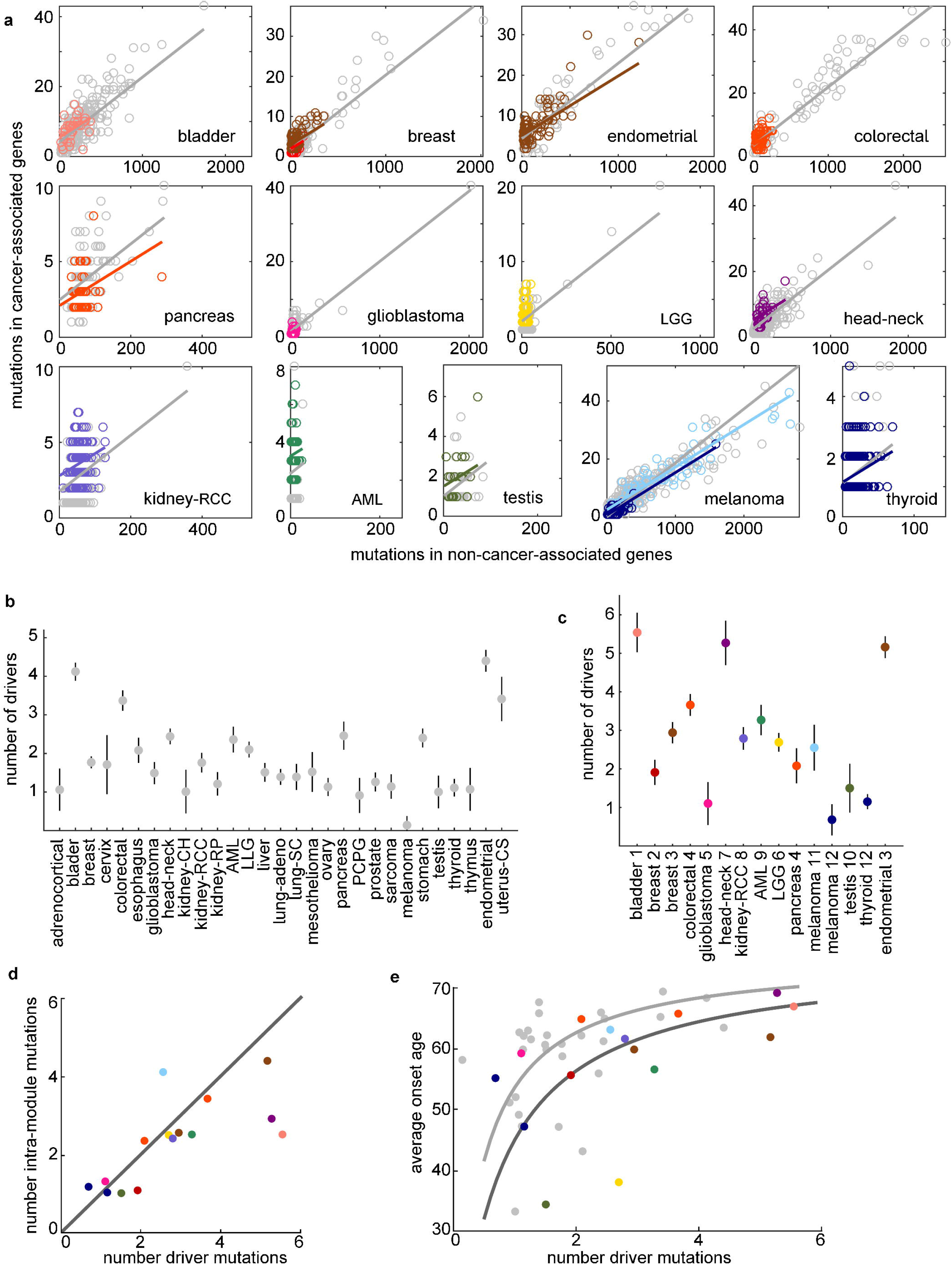
Estimation of the average number of driver mutations per tumor. **(a)** Regression between the number of coding mutations in cancer-associated (y-axis) and non-cancer-associated (x-axis) genes. Colored circles correspond to samples from significant modules. The solid lines show the fit to the ANCOVA model *y* = (*α* + *α_i_*) + *βx* + *ϵ* when considering all samples (gray), or samples from significant modules (colored). All colored lines have identical slope, but differ in their intercept, and the same holds for gray lines. The values of the slope are 0.0186 (gray) and 0.0147 (colored), with global R^2^ = 0.75 and 0.52, respectively (*p* < 10^−20^ in both cases). The intercepts, that correspond to the estimated number of driver mutations, are represented in **(b)** (all cancer types) and **(c)** (members of significant modules); error bars represent 95% confidence intervals. The number of drivers correlates with the number of intra-module mutations **(d)** (Spearman’s rho = 0.772, *p* < 0.001) and with the age at diagnosis **(e)** (Spearman’s rho = 0.527, p = 0.003). Solid lines in **(e)** are fits to the curve 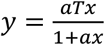 derived from the model of Armitage and Doll19, where *T* = 75 is the average lifespan in the absence of cancer and *a* is the proportionality constant between the number of drivers and rate-limiting steps (light gray, *a* = 2.5, all tumors; dark gray, *a* = 1.5, tumors from significant modules).

Overall, both the number of drivers and the fraction of mutations that are inferred to be drivers tend to be larger in samples from the significant, specific modules. Moreover, the estimated number of driver mutations in these samples closely matches the average number of module-specific (i.e. intra-module) mutations per tumor (Fig. 4D, Spearman’s rho = 0.772, *p* < 0.001). These two findings indicate that modules are held together by genes that carry actual driver mutations in the cancer types that belong to the respective modules. Deviations from the 1:1 trend in Fig. 4D reveal three exceptional cases of limited correspondence between the number of module-specific mutations and the number of estimated drivers. First, the average number of module-specific mutations in bladder and head and neck cancers notably falls below the estimated number of drivers, suggesting the existence of driver mutations in genes that do not belong to the module. Second, the number of module-specific mutations in the high-mutation melanoma module is larger than the number of drivers, suggesting that some of the mutations in genes from this module are passengers.

There is a strong positive correlation between the estimated number of drivers in a cancer type and the average age at which the respective cancers are diagnosed (Fig. 4E). This result holds regardless of whether the number of drivers is estimated for the complete set of samples of the given cancer type (Spearman’s rho = 0.527, p = 0.003) or for the samples assigned to significant modules only (rho = 0.693, p = 0.005). After controlling for the differences in the number of drivers, cancer types that are and are not associated with significant modules (i.e. with and without specific sets of cancer-associated genes) significantly differ in the average diagnosis age, the former appearing earlier in life (ANCOVA on rank-transformed data, p = 0.036, difference = 6.7 rank units).

## Discussion

It is a well-established fact that tumors accumulate recurrent mutations in some genes more often than in others. This phenomenon underlies the discovery of oncogenes and tumor suppressor genes and has led to the identification of central pathways for tumorigenesis and tumor progression^7, 8, 10, 12, 41^. Here we went a step further and tested to what extent the association between genes and cancer types is mutually specific and suffices to define coherent, biologically meaningful groups of tumors. Although related to previous research on detection of significantly mutated genes and tumor classification^26, 42, 43^, our approach differed in three major aspects. First, tumor samples were not classified a priori based on their histology, which enabled us to test if different cancer types are distinguishable through comparison of their mutational landscapes alone. Second, genes and samples are jointly clustered in a single step, so that the resulting network modules reflect mutual specificity. Third, because our network-based clustering is conducive to rigorous statistical testing, we could discriminate between cancer types that do and do not show a significant degree of specificity in their sets of mutated genes.

Previous studies involving 12 major cancer types have shown that tissue-specific clusters can be automatically identified from genomic and transcriptomic features, suggesting the existence of a consistent molecular basis for a tissue-based classification of tumors^26^. Our analysis provides a generalization of that result to a more diverse dataset that included 30 cancer types and 198 cancer-associated genes, revealing major differences among cancer types. Thus, colorectal, pancreatic, endometrial, kidney (clear cell), breast, thyroid, and brain cancers, acute myeloid leukemia, sarcoma, mesothelioma, melanoma and uterine carcinosarcoma are significantly associated with mutations in tissue-specific sets of genes. In contrast, stomach, esophagus and lung cancers, among others, follow a more diffuse, less specific mode of driver accumulation. Some cancers, such as bladder, prostate, testicular (germ cell), and head-and-neck squamous cancer, show a mixed picture, with a significant specificity of mutations observed only in a fraction of samples.

The observed specificity patterns of cancer-associated genes could originate from at least two, not mutually exclusive, causes. Biases in mutation and/or repair could make some tissues more prone to accumulating mutations in certain genes (e.g. due to differences in transcription levels, chromatin configuration and exposure to mutational processes) although such biases are unlikely to account for the large differences observed across cancer types^44^. A more important factor is the tissue-specificity of the pathways that control cell proliferation, which have to be overcome for tumor progression through mutations in different genes^45^. This view is supported by experimental research on the functional mechanisms by which APC and KRAS mutations lead to colorectal cancer^46, 47^ and combined VHL-BAP1 mutations lead to clear-cell renal cell carcinoma^48^. Along similar lines, recent analysis of synonymous and non-synonymous substitutions in cancer genomes has shown that positive selection promotes fixation of somatic mutations in a gene-and-tissue-specific manner, implying that selection pressures during tumorigenesis are tissue-specific^9^.

The absence of detectable specificity patterns in some cancers might be affected by inherent limitations of community detection algorithms on large, partially modular networks, such as the one we analyzed here. In particular, small sample size could compromise the identification of modules for thymoma, adrenocortical, cervical, and kidney (chromophobe) carcinomas. Additionally, it could be hard to find specificity patterns in cancers with high mutational load, such as lung and stomach cancer, due to their low signal (drivers) to noise (passengers) ratio. Nonetheless, the detection of significant modules for bladder cancer and melanoma, which both have high mutational loads^3, 49^, implies that a high mutation burden does not critically affect the performance of the module detection algorithm. Finally, relevant specificity modules based on copy number variation or gene rearrangements could remain undetected if these large-scale mutations involve genes that are not considered here. It should be noted that the absence of specific sets of driver genes for some cancer types does not imply that such cancers cannot be clustered on the basis of other molecular features, as it has been shown for lung adenocarcinoma and lung squamous carcinoma based on transcriptomics^26^.

The second major theme of this study is the estimation of the number of driver mutations affecting cancer-associated genes in different cancer types. By comparing the number of mutations in cancer-associated and other genes, we inferred an average of approximately 2 driver mutations in cancer genes per tumor, with significant variation (from <1 to >5) across cancer types. Our results generally agree with previously reported numbers based of mutations under positive selection, providing an independent support for such values^9^. Remarkably, there is a connection between driver mutations and tissue-specific, cancer-associated genes. Specifically, the number of driver mutations in different cancer types shows a 1:1 correspondence (some minor variations notwithstanding) with the number of intramodule mutations, that is, with the number of mutations in tissue-specific genes.

We further show that the number of driver mutations strongly and positively correlates with the mean age of cancer onset, as one would expect if the number of driver mutations was proportional (yet not necessarily equal) to the number of rate-limiting steps in tumorigenesis^19, 21^. Supporting this view, many of the tissue-specific, cancer-associated genes detected in this study are targets of mutations that appear as early clonal events in the trunk of single-tumor evolutionary trees and likely reflect crucial steps in tumor progression^50^. That is, for example, the case of VHL and PBRM1 in clear-cell renal cell carcinoma (often accompanied by parallel subclonal mutations in SETD2)^51^; DNMT3A and NPM1 in acute myeloid leukemia (often with parallel subclonal mutations in FLT3)^52^; KRAS, TP53 and SMAD4 in pancreatic cancer^53^; and APC, KRAS and TP53 in colorectal cancer^54^.

We also found an intriguing link between cancer onset and the specificity of cancer-associated genes: cancer types that carry mutations in specific genes tend to appear earlier in life than expected given their estimated number of driver mutations. We suspect that this difference could be explained by the requirement for additional driver mutations in genes that are currently not classified as cancer-associated in the case of the non-specific cancer types and/or by stronger effects of driver mutations in the modular group of cancers.

The results of this work show that rigorous statistical methods for community detection in bipartite networks can shed light on the relationships between different types of tumors and cancer-associated genes. The modularity of the gene-cancer network analyzed here is relatively low, with only about one in four tumor samples included in significant modules. This fraction is likely to increase as more tumors are sequenced and additional cancer-associated genes are identified. Nevertheless, the good agreement between the numbers of driver mutations estimated here and those obtained by other methods, as well as the significant difference in the cancer onset age between the modules and the diffuse component of the network, suggest that the difference between the two modes of carcinogenesis revealed by this analysis withstands the test of time. This distinction between tumors caused by driver mutations in limited sets of tissue-specific genes and those caused by mutations in interchangeable genes that are only weakly linked to specific tumor types could have important implications for understanding cancer evolution as well as diagnostic and therapeutic approaches.

## Methods

### Data: tumors and mutations

TCGA public mutation calls were downloaded from the tcga-data.nci.nih.govftp site on January 2016. Mutations were reannotated with the Ensembl Variant Effect Predictor (VEP) software^55^; information on the affected gene, type of mutation and coarse-grained impact was collected. The classification of genes into oncogenes and tumor suppressors was extracted from Ref ^56^.

### Network construction

An unweighted, undirected, bipartite network of somatic mutations in cancers was built by connecting tumor sample nodes to gene nodes whenever a small mutation in the coding region was identified in a given gene in a given sample. For this purpose, the following were considered as coding mutations: missense and nonsense substitutions, loss of a stop codon, mutations affecting splice donor or acceptor sites, in-frame and out-of-frame indels. To keep the network size tractable and maximize the signal-to-noise ratio, only genes with a known association with cancer were included in the network. Specifically, we used the list of 198 cancer genes reported in Ref ^3^, which includes the curated list of 174 mutated genes from the COSMIC database (version 73)^28^ and any other gene from the Cancer Gene Census database found recurrently mutated in Ref ^7^. Samples without any mutation in cancer genes and samples with >3000 coding mutations (hypermutators) were excluded from the network. The resulting network consisted of a single connected component with 7665 samples and 198 genes, with a density of connections of 0.019.

### Module detection

Following recent methodological advances in network analysis, a consensus clustering approach was used to identify the modules of the network^57^. In the first step, maximization of Barber’s modularity index was performed in 200 replicas of the network with the software MODULAR (simulated annealing algorithm, default parameters)^58^, which yielded 200 alternative partitions. To test, from a global perspective, if the cancer mutation network has a significant modular structure, we generated 200 random bipartite networks with the same gene- and sample-degree distributions, ran MODULAR on them, and compared the modularities of the resulting partitions with those of the cancer mutation network, using a Welch’s T-test. In the second step, the 200 alternative partitions of the cancer mutation network were used to build a consensus matrix by assigning to each pair of nodes a value equal to the fraction of replicas in which both nodes were assigned to the same module. A distance matrix was then defined as one minus the consensus matrix, and the consensus partition was finally obtained by performing hierarchical clustering on the distance matrix (UPGMA method, implemented by the ‘linkage’ function in MATLAB version R2015a). The number of clusters was chosen to maximize the Barber’s modularity index of the consensus partition with respect to the original network. We refer to the clusters in this consensus partition as the “unfiltered modules”.

To evaluate the significance of each module separately and filter out the genes and samples that do not follow a modular pattern, we ran the software OSLOM on the original network with the options -singlet -r 0 -hr 0 -t 0.05 -hint, using the unfiltered modules as the reference partition^38^. The significance threshold was set to 0.05. With these settings, OSLOM evaluates the probability that nodes from a random (non-modular) network display the connection patterns observed in the reference partition and removes those nodes that do not reach the required significance threshold. The result is a filtered partition that only includes nodes (genes and samples) with a statistically significant modular structure. As an additional output, OSLOM returns an “extended” partition in which the nodes that do not reach the significance threshold are reassigned to the module that minimizes their p-value. We refer to the modules in such partition as the “extended modules”.

To test whether the results were robust to the choice of the network partitioning method, we repeated the network analysis using Infomap as the module detection software^59^, both in the first and second steps of the consensus clustering pipeline (in this case, all entries with values >0.4 in the consensus matrix were set to 0, and Infomap was run on the network defined by the consensus matrix constructed with this threshold). Both the unipartite and bipartite versions of Infomap (options set to find hard partitions with 2 levels of hierarchy) were run, followed by the assessment of module significance with OSLOM. In all cases, the results were similar to those presented here (Supplementary Table S8).

### Estimation of the number of driver mutations

To estimate the number of driver mutations in different cancer types, we considered all small coding mutations affecting cancer-associated and non-cancer-associated genes. The function ‘aoctool’ in MATLAB R2015a was used to fit the number of mutations in cancer genes to a “parallel lines” linear model with cancer type and number of mutations in non-cancer-associated genes as independent variables. The statistical significance of the model parameters was evaluated with an ANCOVA, with cancer types as factors and the number of mutations in non-cancer-associated genes as covariable. The entire analysis was carried out twice. First, all the samples in the dataset, classified into cancer types, were analyzed. Then, the analysis was limited to the samples that were represented in any of the significant modules. Under the second approach, cancer types represented by fewer than 20 samples were excluded, and cancer types assigned to more than one significant module were split to obtain module-specific estimates.

The association between the number of drivers and the age of diagnosis was initially evaluated through a Spearman’s correlation analysis. To further assess whether cancer types associated or not to significant modules differ in their age of diagnosis while controlling for the number of driver mutations, an ANCOVA was performed on the rank-converted data, with the number of drivers as a covariable and the inclusion into a significant module as a binary factor.

## Acknowledgements

J.I. and E.V.K. are supported by intramural funds of the US Department of Health and Human Services (to the National Library of Medicine). I.M. is supported by EMBO (aALTF 766-2015) and Cancer Research UK (C57387/A21777).

## Author contributions

J.I. and E.V.K conceived the study; I.M. downloaded the TCGA data and provided key advice; J.I. carried out the analysis; J.I., I.M., and E.V.K. interpreted the results; J.I. and E.V.K wrote the manuscript.

